# Retroelement decay by the exonuclease XRN1 is a viral mimicry dependency in cancer

**DOI:** 10.1101/2023.03.30.531699

**Authors:** Amir Hosseini, Håvard T. Lindholm, Raymond Chen, Parinaz Mehdipour, Sajid A. Marhon, Charles A. Ishak, Daniel D. De Carvalho

## Abstract

Viral mimicry describes the immune response induced by endogenous stimuli such as dsRNA formed by endogenous retroelements. Activation of viral mimicry has the potential to kill cancer cells or augment anti-tumor immune response. Paradoxically, cancer cells frequently present a dysregulated epigenome, leading to increased expression of retroelements. We previously found that ADAR1 p150 upregulation is an adaptation mechanism to tolerate high retroelement-derived dsRNA levels, leading to a druggable dependency. Here, we systematically identified novel mechanisms of viral mimicry adaptation associated with cancer cell dependencies. We correlated the gene knockout sensitivity from the DepMap dataset and interferon stimulated gene (ISG) expression in the Cancer Cell Line Encyclopedia (CCLE) dataset of 1005 human cell lines and identified pathways such as RNA modification and nucleic acid metabolism. Among the top hits was the RNA decay protein XRN1 as an essential gene for the survival of a subset of cancer cell lines. XRN1-sensitive cancer cell lines have a high level of cytosolic dsRNA and high ISG expression. Furthermore, sensitivity to XRN1 knockout was mediated by MAVS and PKR activation, indicating that the cells die due to XRN1-dependent induction of viral mimicry. XRN1-resistant cell lines had low basal dsRNA levels, but became synthetically dependent on XRN1 upon treatment with viral mimicry inducing drugs such as 5-AZA-CdR or palbociclib. Finally, XRN1-dependency is partly independent of ADAR1 activity. These results confirm the potential for our ISG correlation analysis to discover novel regulators of viral mimicry and show that XRN1 activation is an adaptive mechanism to control high dsRNA stress induced by dysregulated retroelements in cancer cells and creates a dependency that can be explored for novel cancer therapies.

## Introduction

When a human cell detects a viral infection, it enters an antiviral state with tailored responses to defeat the infection. For example, the cell will produce interferons (IFNs) that reduce protein synthesis to hinder viral replication, increase presentation of peptides to the immune system, induce apoptosis, and signal in a paracrine manner to stimulate the same antiviral response in neighboring cells^1^. Interestingly, this antiviral state can also be activated from endogenous stimuli, such as double-stranded RNA (dsRNA) structures derived from endogenous retroelements, in a phenomenon called ‘viral mimicry’^2–4^. Activation of viral mimicry is a promising approach for cancer therapeutics because in the same way the antiviral response can help the body eliminate cells infected with viruses, it can be leveraged to kill cancer cells^5–8^.

Endogenous retroelements are sections of DNA that can duplicate within the genome through an RNA intermediate^9^. In homeostasis, endogenous retroelements are typically repressed through a variety of mechanisms in order to prevent deleterious effects such as genomic instability or inappropriate immune activation. Therapeutic disruption of these mechanisms, such as with the DNA demethylating agent 5-aza-2’-deoxycytidine (5-AZA-CdR), can lead to retroelement expression and formation of dsRNA structures which bind to innate immune receptors and produce a viral mimicry response^3–5^. Beyond DNA methylation, disruption of histone modifications^6,10^, splicing^7,11^, and RNA editing^5,12–14^ have also been implicated in viral mimicry-mediated anti-cancer effects. Due to the wide variety of cellular processes involved, we have proposed that in addition to their role in cancer treatment, endogenous retroelements can act as an alarm for disruptions to cellular homeostasis that culls pre-cancerous cells^15^. Despite this, cancers often paradoxically display elevated levels of endogenous retroelements^2^, suggesting that these cancers have developed mechanisms of viral mimicry adaptation. Such mechanisms may represent novel cancer dependencies that can be targeted therapeutically to re-sensitive tumors to viral mimicry.

Previously, we identified upregulation of the p150 isoform of the RNA editing enzyme adenosine deaminase acting on RNA 1 (ADAR1) as one such adaptation^5^. ADAR1 catalyzes adenosine-to-inosine (A-to-I) editing primarily in RNA duplex structures derived from Alus^16^, a family of endogenous retroelements that make up about 10% of the human genome^17^. Adjacent Alus found on the same strand but in opposite orientations are known as inverted-repeat Alus (IR-Alus) and can form dsRNA hairpin structures when transcribed together^5^. In the absence of ADAR1, IR-Alu dsRNAs are recognized by innate immune receptors such as melanoma differentiation-associated protein 5 (MDA5)^18^, which then activates IFN signaling through the mitochondrial antiviral-signaling protein (MAVS) pathway. The subsequent expression of interferon-stimulated genes (ISGs) results in an antiviral cell state. A-to-I editing by ADAR1 creates bulges in the dsRNA structure that prevent MDA5 recognition^18^, allowing cancer cells with upregulated ADAR1 to circumvent viral mimicry activation. However, this ADAR1 dependency provides a novel vulnerability that can be exploited for cancer therapy. Accordingly, our previous work indicates that depletion of ADAR1 potentiates viral mimicry-inducing epigenetic therapies^5^.

Here, we systematically screened for other novel viral mimicry dependencies by correlating ISG expression in cell lines from the Cancer Cell Line Encyclopedia (CCLE) with cell viability after genetic knockout as determined in the DepMap dataset^19,20^. This analysis identified pathways of interest, including RNA modification and nucleic acid metabolism pathways. Among the top hits for individual genes was the RNA decay protein XRN1, which degrades RNA without a 5’ cap in the 5’ to 3’ direction^21^. XRN1 dependency was associated with high basal levels of viral mimicry activation, as determined by dsRNA levels and ISG expression, and was mediated by MAVS and protein kinase R (PKR). Furthermore, while cancer cells with low baseline viral mimicry levels were XRN1-independent, we were able to create a synthetic XRN1 dependency through treatment with drugs that increase dsRNA levels. These results highlight XRN1 as a promising therapeutic target partly independent of ADAR1 and demonstrate the utility of our screening approach, suggesting that validation of other identified hits may reveal additional viral mimicry dependencies.

## Results

### Identification of genes regulating viral mimicry adaptation

Disruptions to cellular mechanisms are common in cancer and can lead to the presence of immunogenic endogenous retroelements and viral mimicry^15^. To survive this viral mimicry induction, cancer cells depend on adaptation mechanisms such as A-to-I editing of dsRNA by ADAR1 to avoid viral mimicry-induced cell death^5^. To broadly define viral mimicry activation at baseline in different cancer types, we calculated the average scaled ISG expression for 1005 human cell lines in the CCLE using a previously published definition of ISG expression (Supplementary Table 1)^22^. We reasoned that the cell lines with high ISG expression would depend on viral mimicry-inhibiting genes such as ADAR1 to adapt to the high levels of viral mimicry. To test this hypothesis, we compared ISG induction to the effect of ADAR1 knockout on viability from the DepMap CRISPR dataset in 1005 cell lines (Fig. 1A)^20^. We found a significant correlation between ADAR1 knockout-mediated decrease in viability and ISG expression across the cell lines, suggesting that cell lines with high ISG expression depend on ADAR1 to survive the induction of viral mimicry, confirming previous findings^5,22,23^. To search for novel viral mimicry adaptation mechanisms, we extended the analysis to the correlation between ISG expression and viability upon individual knockout of all genes (Fig. 1B). This analysis revealed genes that cancer cells depend on to survive viral mimicry induction and are therefore potential targets for cancer therapeutics. Among the hits, ADAR1 had the highest correlation between ISG expression and effect on viability.

**Figure 1.**
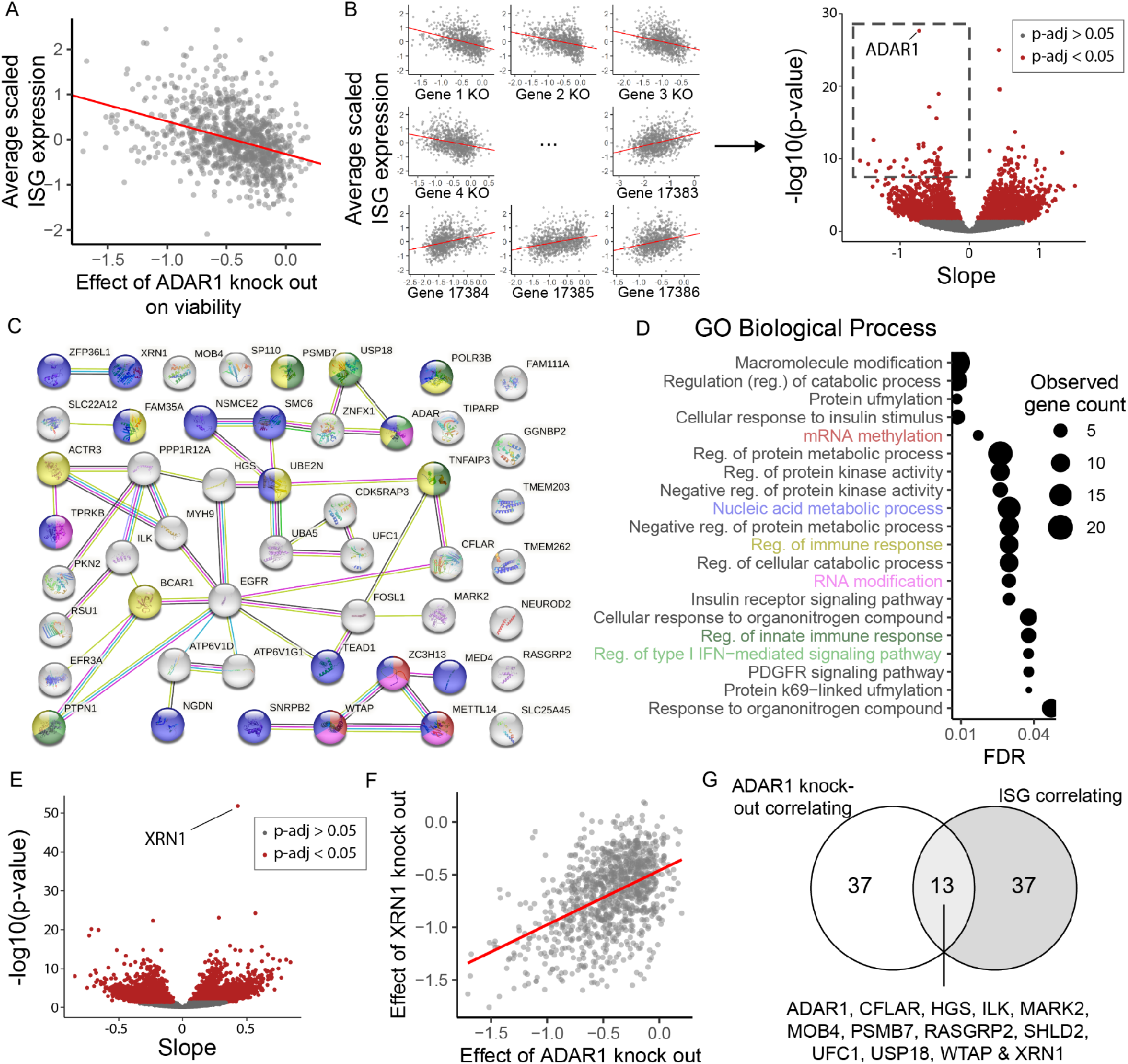
Discovery of genes regulating viral mimicry adaptation. (A) Linear correlation between effect of ADAR1 knockout on viability from the CRISPR knockout dataset from DepMap and the average scaled expression of ISGs from CCLE. Each dot represents a cell line, n = 1005. The value on the x-axis is the CERES score from DepMap representing the effect size on viability from knocking out ADAR, normalized against a distribution of non-essential and pan-essential genes. The value on the y-axis is the mean of z-score normalized log(TPM+1) values from 38 ISGs defined in Liu et al.^22^ Negative value is more sensitive to ADAR1 knockout. R = -0.36, Benjamini-Hochberg corrected p-value for linear correlation = 2.4 × 10^−28^ (B) Analysis in Figure 1A extended to all genes with the resulting slopes and Benjamini-Hochberg corrected p-value for each gene plotted in a volcano plot. The red square represents the top 50 most significant genes with negative slope. (C) Network from the STRING tool showing the top 50 most significant genes with negative slope (area outlined in B). (D) Enrichment analysis with the STRING tool showing the 20 GO biological process terms with the highest strength value as determined by STRING. The color of GO term names corresponds to the node color seen in C. (E) Slope and Benjamini-Hochberg corrected p-value from the linear correlation between the effect on viability from knocking out ADAR1 from the CRISPR knockout DepMap dataset and all other genes. (F) Linear correlation between the viability effects of XRN1 knockout and ADAR1 knockout in the CRISPR knockout DepMap dataset. R = 0.47, Benjamini-Hochberg corrected p-value = 1.8 × 10^−52^ (G) Comparing the top 50 genes by p-value with negative slope in B with the top 50 genes with positive slope in E.

To explore novel mechanisms of viral mimicry adaptation, we selected the top 50 genes with negative slope and clustered them with the STRING tool (Fig. 1C)^24^. Among these 50 genes we found enrichment of immune activation pathways such as “Regulation of immune response” and “Regulation of type I IFN-mediated signaling pathway” (Fig. 1D). As dsRNA is a common activator of viral mimicry, it is interesting that we found GO terms related to RNA processing such as “mRNA methylation” and “nucleic acid metabolic process”. Disruption of RNA methylation has been previously linked to innate immune activation from endogenous retroelements^12–14^.

Since high ISG expression can occur through multiple mechanisms, the correlation between high ISG expression and dependency to a gene could reveal diverse mechanisms for viral mimicry adaptation. To focus our analysis on targets that regulate the presence of immunogenic endogenous retroelements we also compared the dependency of a gene to the sensitivity to ADAR1 knockout. ADAR1 was the top hit correlating with ISG expression and has been previously shown to inhibit the activation of innate immune receptors by dsRNA from endogenous retroelements and thus viral mimicry^5^. We therefore did a linear regression analysis between sensitivity to ADAR1 knockout and the sensitivity to knockout of other genes and plotted the slope and p-value for each correlation (Fig. 1E). Out of all genes, the RNA decay protein XRN1 had the most significant correlation with ADAR1 (Fig. 1E and 1F). XRN1 was also among the top 50 most correlating genes in both the ISG correlation analysis and ADAR1 knockout correlation analysis (Fig. 1G). Interestingly, when we correlated the effect on viability of XRN1 knockout with gene expression for all genes, we found that the most highly correlated genes are enriched with pathways connected to an antiviral state (Fig. S1A-D). Out of the top 50 genes, 31 are part of the GO term “Defense response to virus” (Fig. S1C-D). This analysis demonstrates a clear connection between baseline ISG induction and sensitivity to XRN1 depletion.

### XRN1 dependency in cancer cell lines is associated with elevated ISG expression and dsRNA levels

We reasoned that cells with higher baseline dsRNA accumulation would be more dependent on RNA decay proteins such as XRN1. To explore the mechanism of XRN1 dependent adaptation to viral mimicry induction, we used the DepMap dataset to classify cell lines as either XRN-sensitive (had lower viability after XRN1 knockout) or XRN-resistant (viability was not affected by XRN1 knockout). We selected two colorectal cancer cell lines (C84 and NCI-H747) and two breast cancer cell lines (HCC202 and MDA-MB-468) that are XRN1-sensitive and two colorectal cancer (COLO678 and NCI-H716) and two breast cancer cell lines (EFM19 and MCF7) that are XRN1-resistant (Fig. S1E). To further explore our findings in clinically relevant models, we also included the patient-derived colon cancer stem cell (CSC)-enriched spheroid line POP92^25,26^. In order to validate the results from our DepMap analysis, we knocked out XRN1 in these models with CRISPR-Cas9 (Fig. S2A). The XRN1-sensitive cell lines NCI-H747 and C84 had highly reduced numbers five days after XRN1 knockout compared to the XRN1-resistant cell lines NCI-H716 and COLO678 (Fig. 2A, 2B and S2B). Additionally, we found that the XRN1-sensitive cell line NCI-H747 had an elevated expression of ISGs at baseline compared to the XRN1-resistant cell lines COLO678 and NCI-H716 (Fig. 2C). POP92 cells have low ISG expression and are not sensitive to XRN1 knockout, categorizing it as XRN1-resistant (Fig. 2A, 2C).

**Figure 2.**
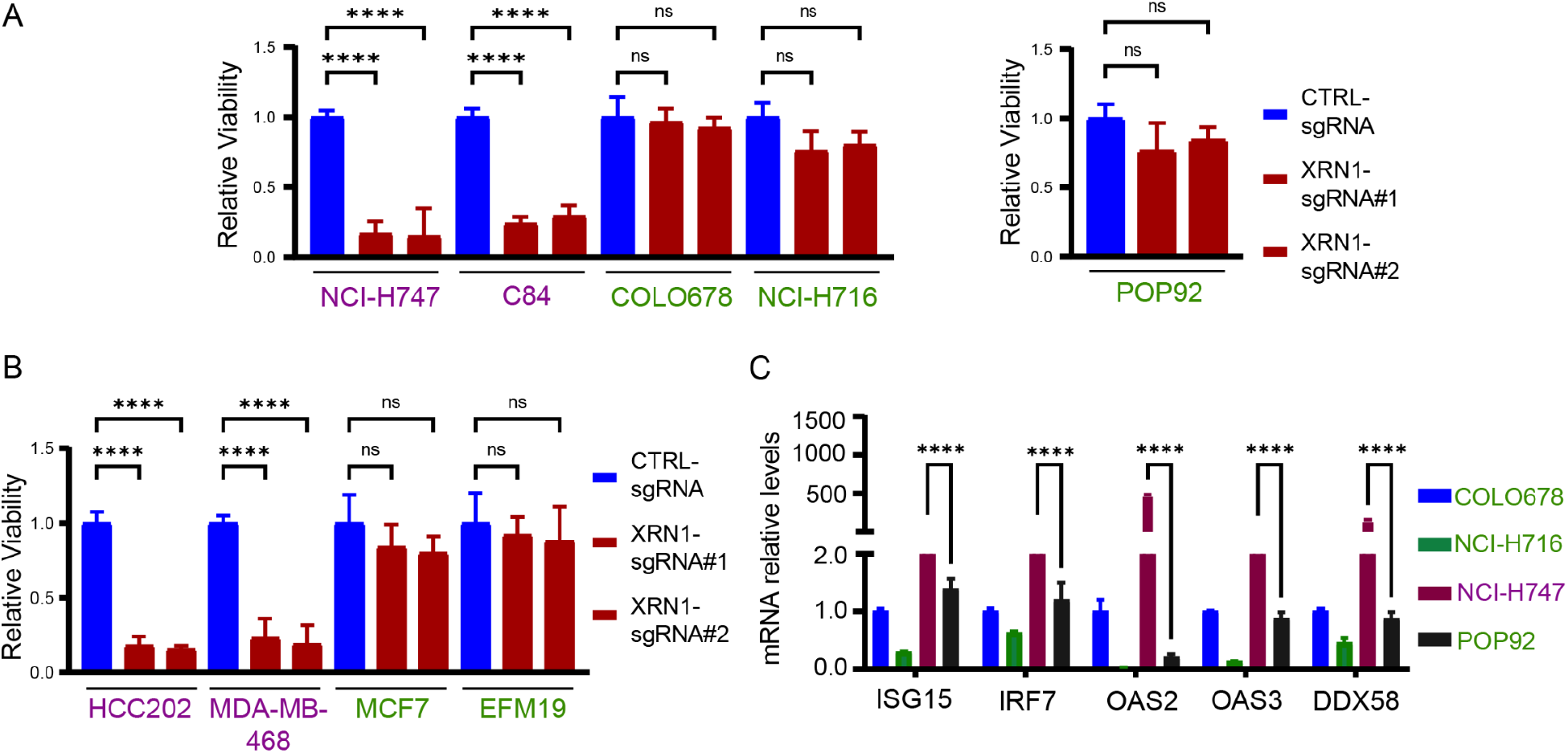
A subset of cancer cell lines are sensitive to XRN1 knockout. (A, B) Cell viability assessed by CellTiter-Glo, 5 days after XRN1 knockout with CRISPR-Cas9 in colorectal (A) and breast (B) cancer cell lines. XRN1-sensitive cell lines are colored purple and XRN1-resistant cell lines are colored green. Data presented as the mean of duplicates for C84 and triplicates for other cell lines ± SD. (C) qPCR of indicated ISGs in colorectal cancer cells. Values are normalized against RPLP0. Data presented as the mean of triplicates ± SD.

Next, we sought to determine whether elevated levels of ISGs in XRN1-sensitive cells are due to high baseline levels of cytosolic dsRNAs. Dot blots of total RNA using the J2 antibody, which recognizes dsRNA helixes longer than 40 base pairs, revealed elevated dsRNA levels in XRN1-sensitive cell lines compared to XRN1-resistant cell lines (Fig. 3A). Confocal imaging staining for dsRNA with the J2 antibody confirmed the increased presence of cytosolic dsRNA in the XRN1-sensitive cell line NCI-H747 compared to the XRN1-resistant cell lines COLO678 and NCI-H716 (Fig. 3B). Collectively, XRN1 dependency is associated with elevated ISG expression and a high level of cytosolic dsRNA, which are both hallmarks of viral mimicry induction.

**Figure 3.**
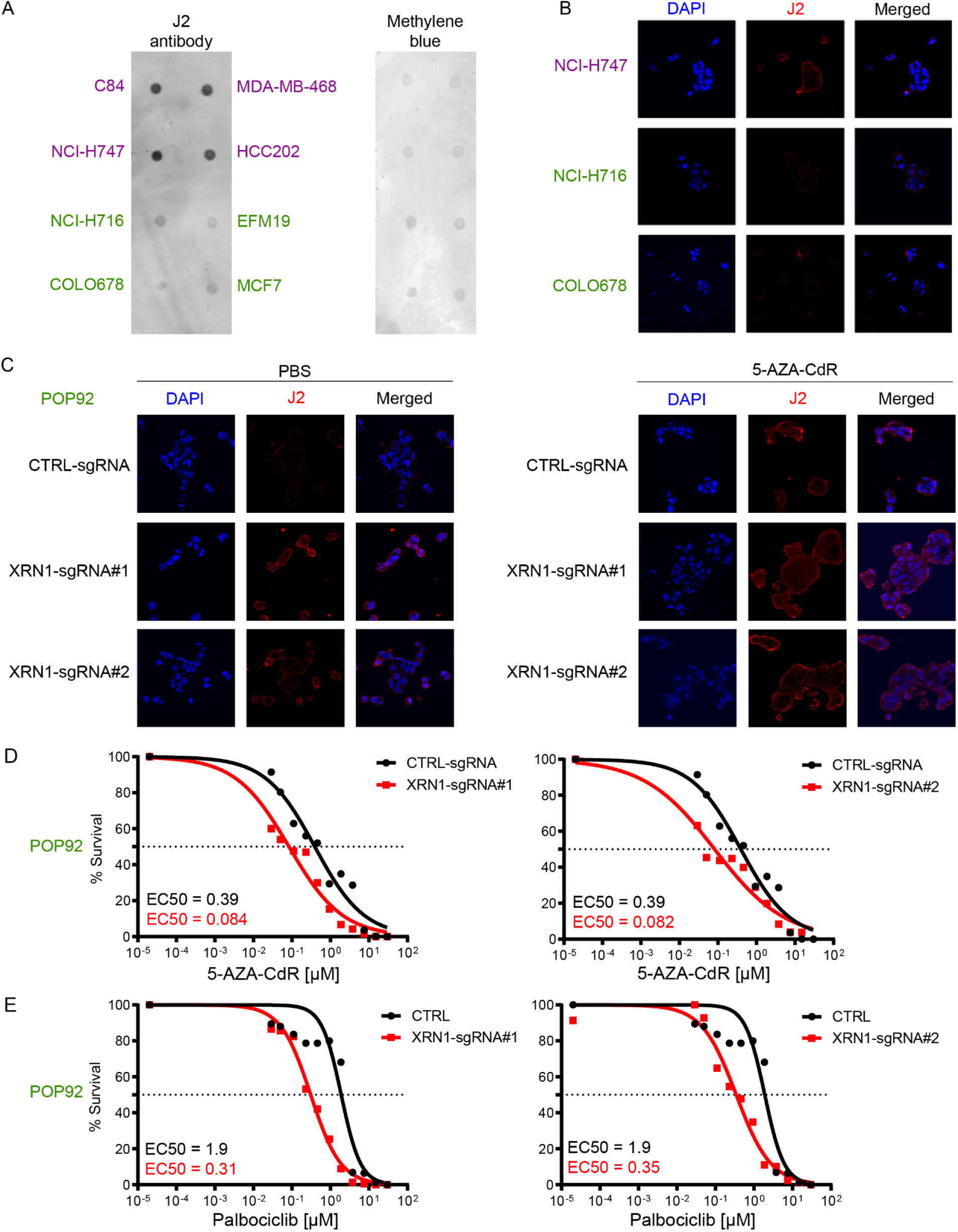
dsRNA induced by Palbociclib or 5-AZA-CdR produces a synthetic dependency to XRN1 in XRN1-resistant POP92 cells. (A) Dot blot for dsRNA using total RNA from indicated cell lines. Normalized amounts of total RNA were dotted on Hybond N+ membranes, visualized by methylene blue staining, and immunoblotted with J2 antibody. (B) Representative confocal microscopy images of colorectal cell lines. Nuclei were stained with DAPI (blue) and dsRNA was stained using the J2 antibody (red). (C) Representative confocal microscopy images from control and knockout of XRN1 of POP92 cells treated with PBS or 5-AZA-CdR. Nuclei were stained with DAPI (blue) and dsRNA was stained using the J2 antibody (red). (D) Survival of wild-type XRN1 (black) and XRN1 knockout (red) patient-derived CRC cells (POP92) after treatment with 5-AZA-CdR. Luminescence signal was normalized, and dose-response curves and EC50 values were calculated using a nonlinear regression curve fit. (E) Survival of wild-type XRN1 (black) and XRN1-knockout (red) POP92 cells after treatment with palbociclib. Luminescent signal was normalized, and dose-response curves and EC50 values were calculated using a nonlinear regression curve fit.

### Pharmacological induction of cytosolic dsRNA produces a synthetic dependency on XRN1

We next sought to determine whether increased levels of cytosolic dsRNA could sensitize XRN1-resistant cells to XRN1 inhibition. DNA demethylating agents such as 5-AZA-CdR and CDK4/6 inhibitors such as palbociclib can induce viral mimicry by increasing the level of cytosolic dsRNA^3,4,27^. Immunostaining in POP92 XRN1 knockout cells with the J2 antibody revealed a significant increase in cytoplasmic dsRNA in XRN1-depleted cells treated with 5-AZA-CdR compared to control cells (Fig. 3C). XRN1 knockout led to a significant reduction of the 5-AZA-CdR half-maximal effective concentration (EC50) *in vitro* (Fig. 3D).

Furthermore, 5-AZA-CdR sensitized the four other XRN1-resistant cell lines to XRN1 knockout (Fig. S3A). The EC50 of palbociclib was also reduced by over five-fold in POP92 cells with XRN1 knocked out compared to control. Like 5-AZA-CdR, palbociclib induces a synthetic dependency on XRN1 in XRN1-resistant cancer cell lines (Fig. S3B). Collectively, these results suggest that sensitivity to XRN1 may be mediated by cytoplasmic dsRNA levels and that XRN1-resistant cell lines can be sensitized to XRN1 knockout by increasing dsRNA levels with viral mimicry-inducing drugs.

### MAVS and PKR activation mediate sensitivity to XRN1 disruption

Cytosolic dsRNAs can trigger an innate immune response upon recognition by cytosolic RNA-sensing receptors such as MDA5 and PKR. Upon dsRNA recognition, MDA5 promotes aggregation of MAVS on the mitochondrial membrane and activates type I/III IFN signaling to induce the expression of ISGs. PKR and ribonuclease L (RNaseL) exert antiviral activity through translational arrest and RNA degradation, respectively^28,29^. To investigate the mechanism of cell lethality after depletion of XRN1, we knocked out MAVS, PKR, and RNaseL (Fig. 4A). In the XRN1-sensitive cell line NCI-H747, we found that both MAVS knockout and PKR knockout partially rescued cell lethality after XRN1 knockout (Fig. 4B and C). Double knockout of MAVS and PKR was able to almost completely rescue the loss of viability in NCI-H747 cells with knockout of XRN1, indicating that these are the major pathways involved in mediating the observed cell viability effects of XRN1 depletion. However, deletion of RNaseL could not rescue cell lethality, indicating that RNaseL is dispensable for this phenotype. Although both MAVS and PKR knockout can affect *ISG15* expression after XRN1 knockout in NCI-H747 cells, we find that MAVS knockout decreases *ISG15* expression to a higher degree than PKR knockout (Fig. 4D). Loss of both PKR and MAVS abrogates XRN1 knockout-induced *ISG15* expression (Fig. 4D).

**Figure 4.**
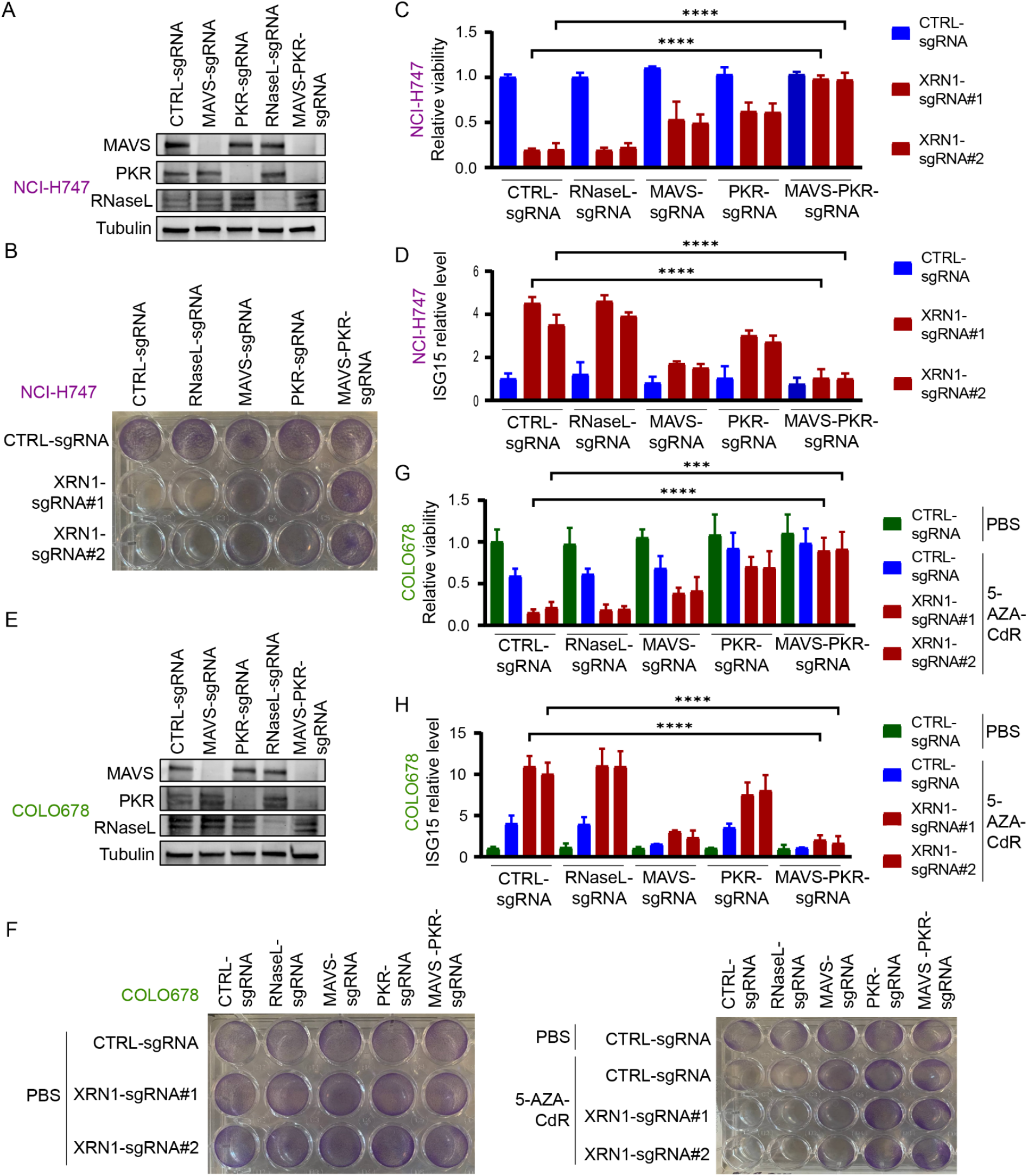
XRN1-dependency requires MAVS signaling and PKR activation. (A) Western blot of MAVS, PKR, RNaseL protein levels in NCI-H747 cells. Tubulin is loading control. (B) Cell viability in NCI-H747 cells by crystal violet staining. (C) Cell viability in NCI-H747 cells by CellTiter-Glo assay. Data presented as the mean of triplicates ± SD. (D) qPCR of *ISG15* in NCI-H747 cells. Values are normalized against *RPLP0*. Data presented as the mean of triplicates ± SD. (E) Western blot of MAVS, PKR, RNaseL protein level in COLO678 cells. Tubulin is loading control. (F) Cell viability in COLO678 cells by crystal violet staining. (G) Cell viability in COLO678 by Cell Titer Glo assay. Data presented as the mean of triplicates ± SD. (H) qPCR of *ISG15* in COLO678 cells. Values are normalized against *RPLP0*. Data presented as the mean of triplicates ± SD.

Since 5-AZA-CdR leads to a synthetic XRN1 dependency in XRN1-resistant cells (Fig. 3D and S3A), we investigated how the loss of MAVS, PKR, and RNaseL modulated the response to 5-AZA-CdR treatment in XRN1 knockout cells (Fig. 4E and F). Similar to NCI-H747 cells, both MAVS and PKR knockout partly rescued the loss of viability upon XRN1 knockout in COLO678 cells treated with 5-AZA-CdR, but double knockout of MAVS and PKR completely rescued the loss in viability (Fig. 4F and G). The degree of rescue observed with PKR knockout in COLO678 cells was more robust compared to cells with MAVS knockout, suggesting that PKR binding of the dsRNA induced by 5-AZA-CdR was mainly responsible for the observed cell death phenotype. On the contrary, MAVS knockout more strongly inhibited *ISG15* induction than PKR knockout cells (Fig. 4H). Altogether, these results suggest that while it is the MAVS pathway that is mainly responsible for ISG activation, both the MAVS and PKR pathways mediate XRN1 knockout-induced cell death. Furthermore, as both of these pathways induce cell death in response to dsRNA, these results highlight the importance of dsRNA in XRN1-dependent cell death.

### XRN1 dependency is partly independent of ADAR1 activity

Cancer cells can avoid cell death induced by viral mimicry through ADAR1-dependent A-to-I editing, which shields dsRNAs from detection by innate immune receptors^5,18^. We therefore investigated whether ADAR1-dependent viral mimicry adaptation can inhibit XRN1 knockout-induced cell death. A-to-I editing levels in these cell lines were estimated by a modified pipeline based on variant calling in RNAseq data from CCLE^19^, and revealed active A-to-I editing with 19,255 and 20,023 A-to-I editing loci in the XRN1-resistant cell lines NCI-H716 and COLO678, respectively, and 24,808 A-to-I editing loci in the XRN1-sensitive cell line NCI-H747 (Fig. 5A). Furthermore, between 30-43% of these A-to-I edits are found in Alus. To investigate the interaction between A-to-I editing and XRN1 knockout-induced viral mimicry, we generated POP92 ADAR1 knockdown cells (Fig. S4A). Although POP92 cells with ADAR1 knockdown have reduced viability upon treatment with 5-AZA-CdR, knockout of XRN1 led to a greater reduction in viability, which was not enhanced by simultaneous XRN1 and ADAR1 depletion (Fig. 5B). Furthermore, we found that ADAR1 overexpression is associated with increased cell viability and lower ISG expression compared to control after treatment with 5-AZA-CdR (Fig. S4B, 5C), confirming that ADAR1 can function as a viral mimicry adaptation mechanism^5^.

**Figure 5.**
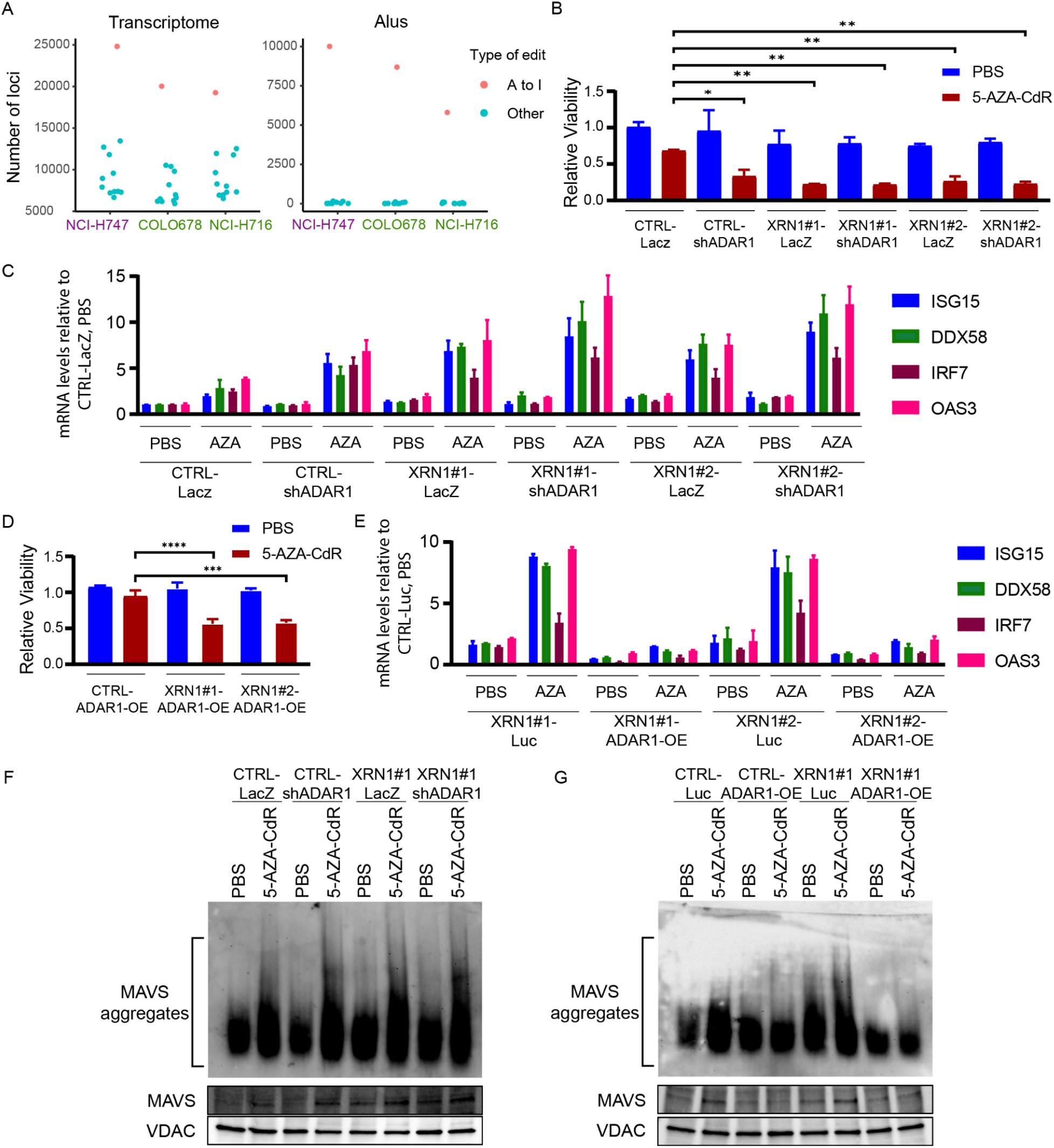
XRN1 depletion can kill cancer cells despite viral mimicry adaptation by high expression of ADAR1. (A) Number of A to I edited loci in the transcriptome and only in Alus in indicated cell lines. Others represent other variants than A to I such as G to C, A to C, etc. (B) Cell viability in POP92 cells with indicated knockout determined with Cell Titer Glo assay. Data presented as the mean of triplicates ± SD. (C) PCR of selected ISGs in POP92 cells with indicated knockout and overexpression of ADAR1 and XRN1. Values are normalized against RPLP0. Data presented as the mean of triplicates ± SD. (D) See B. (E) See C. (F,G) Western blot of MAVS showing aggregation after indicated treatment in POP92 cells with indicated knockout or overexpression of XRN1 and ADAR1. VDAC is loading control.

Interestingly, XRN1 knockout in combination with 5-AZA-CdR in POP92 cells leads to a reduction in cellular viability despite ADAR1-dependent viral mimicry adaptation (Fig 5D). We found higher levels of ISG expression after combined depletion of XRN1 and ADAR1 relative to ADAR1 depletion alone, indicating that XRN1 and ADAR1 together regulate ISG expression from viral mimicry activation (Fig. 5C). However, ADAR1 overexpression was able to prevent ISG induction from treatment with 5-AZA-CdR in cells treated with both control and XRN1 sgRNA (Fig. 5E). Next, we measured aggregation of the mitochondrial protein MAVS in the mitochondria fraction of cell lysates. MAVS aggregation assays confirmed that MAVS aggregates upon treatment with 5-AZA-CdR, and that ADAR1 overexpression was enough to prevent the MAVS aggregation in XRN1 knockout cell lines treated with 5-AZA-CdR (Fig. 5F, G). This suggests that ADAR1 mainly inhibits cell proliferation by interfering with activation of the MDA5/MAVS pathway while XRN1 is active in suppressing both MDA5/MAVS and PKR activation (Fig. 6). These results indicate that targeting XRN1 could have anti-cancer effects that are independent from viral mimicry adaptation by ADAR1 and highlights the therapeutic potential of XRN1.

**Figure 6.**
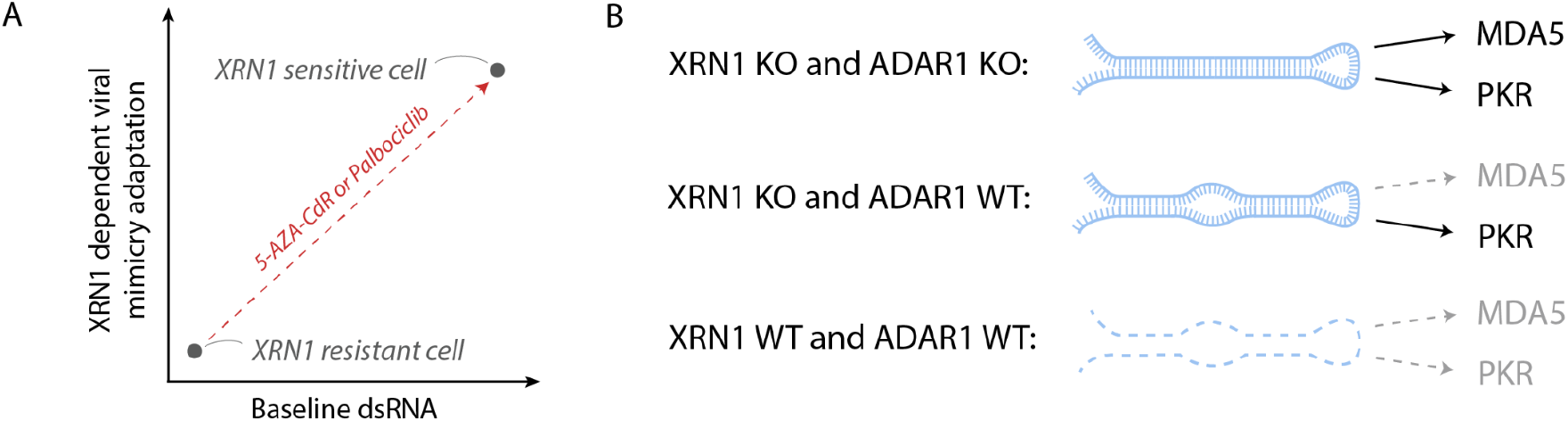
Proposed mechanism for XRN1 dependent viral mimicry adaptation. A) XRN1 resistant cell lines have low levels of immunogenic dsRNA and are therefore not relying on XRN1 to resist viral mimicry induced cell death. XRN1 sensitive cell lines require XRN1 to resist high levels of immunogenic dsRNA to induce viral mimicry induced cell death. Viral mimicry inducing drugs such as 5-AZA-CdR and palbociclib can generate a synthetic dependency to XRN1. B) XRN1 degrades dsRNA while ADAR1 edits A-to-I in dsRNA, these mechanisms have different effects on activation of MDA5 and PKR pathways. When ADAR1 and XRN1 are not present, both MDA5 and PKR can bind dsRNA and activate viral mimicry. If XRN1 is knocked out and only ADAR1 is present, the edited dsRNA will not activate the MDA5 pathway while the PKR pathway can still be activated. If both XRN1 and ADAR1 is present, dsRNA is both edited and degraded which hinders activation of MDA5 and PKR pathways.

## Discussion

To escape the mechanisms limiting cellular growth in normal cells, cancer cells must disrupt normal cellular processes. However, disruption to certain cellular mechanisms regulating expression and RNA metabolism can also lead to the presence of immunogenic endogenous retroelements and induce viral mimicry^15^. To avoid viral mimicry-induced cell death, cancer cells are dependent on viral mimicry adaptation mechanisms such as A-to-I editing of dsRNA by ADAR1^5^. Here we correlated ISG expression, a readout of the antiviral state in the cell, to the effect of gene knockout on viability to survey potential viral mimicry dependencies. Among the top hits we found the RNA decay protein XRN1. Elevated levels of dsRNA and ISG expression were associated with the reduction in viability observed upon XRN1 knockout, which was rescued by double knockout of MAVS and PKR. These are all hallmarks of viral mimicry induction, and confirms that the reduction in viability following XRN1 knockout occurs through induction of viral mimicry.

The most well-described source of endogenous stimuli that can activate viral mimicry are dsRNAs from IR-Alus^5,18^. It is therefore not surprising that proteins associated with RNA processing, such as XRN1, appeared among the viral mimicry adaptation mechanisms appearing in our screen. The most direct explanation for how disrupting XRN1 leads to an antiviral state in the cell is that dsRNA which otherwise would have been degraded by XRN1 accumulates in the cytoplasm, where it can activate the innate immune receptors MDA5 and PKR. Furthermore, our work demonstrates that XRN1-sensitive cell lines already have elevated dsRNA levels as measured by the J2 antibody and thus may be closer to crossing a “threshold” of tolerable dsRNA levels that XRN1 depletion pushes them past. Additional evidence for this mechanism comes from the fact that XRN1-resistant cell lines that have low baseline dsRNA levels can be made sensitive through treatment with dsRNA-elevating drugs like 5-AZA-CdR or palbociclib, suggesting that XRN1 knockout alone is ineffective in these cells because it cannot push their low baseline levels of dsRNA beyond a tolerable threshold. Together, these results indicate that XRN1-dependent degradation of RNA is an important mechanism that prevents accumulation of dsRNA beyond the threshold required for activation of viral mimicry.

Our work further demonstrates that XRN1 knockout-induced cell death is mediated by both MAVS and PKR. We show that MAVS-dependent cell death upon knockout of XRN1 is associated with higher ISG induction, implicating activation of antiviral pathways in this phenotype. PKR-mediated cell death, on the other hand, appears to occur more directly without ISG induction. The fact that PKR and MDA5 are activated by dsRNA of different lengths supports the observation that these two pathways can act independently from each other. PKR can be activated from dsRNA that is as short as 30 bp, while MDA5 preferentially activates the IFN pathway from longer dsRNA molecules^30^. Double knockout of both pathway components was required to generate a near-complete rescue of XRN1 knockout. In contrast, Pestal et al. show that the embryonic lethality induced by depletion of the ADAR1 p150 isoform can be rescued completely by either MDA5 or MAVS knockout alone^31^. We found that ADAR1 overexpression can inhibit XRN1 knockout-induced expression of ISGs, confirming this effect. However, it is interesting that XRN1 knockout reduced cell numbers even when ADAR1 is overexpressed, which suggests that disrupting XRN1 can lead to cell death through activation of PKR pathways in addition to MDA5-MAVS. It is therefore possible that XRN1 is able to evade ADAR1-dependent resistance to viral mimicry induced cell death.

Induction of viral mimicry can be a mechanism for treating cancer either by enhancing cancer cell killing by immune cells or by leading to activation of internal cell death pathways in the cancer cell^2^. Work done in parallel to this study found that silencing XRN1 can potentiate immunotherapy efficiency in mice^32^.

Ran et al. found that disruption of XRN1 activated the IFN pathway through MAVS signaling. Knocking out *Pkr, Sting* or *Mda5* in mice did not completely rescue the ISG induction seen in cells where XRN1 function is disrupted, indicating that XRN1 can induce viral mimicry through multiple pathways.

Interestingly, this study did not find that disrupting XRN1 knockout itself inhibited the cell growth of the mouse tumor cells, indicating that the tumor model they used may be similar to the XRN1-resistant cell lines described here. Our work suggests that disrupting XRN1 alone in an XRN1-resistant tumor would be unlikely to clear the cancer. However, we were able to create a synthetic XRN1 dependency by treating with viral mimicry-inducing drugs such as 5-AZA-CdR and palbociclib. Alternatively, it would be interesting to see if XRN1 can be targeted in tumors that have a high baseline induction of viral mimicry. For example, gliomas with the tumor-driving K27M mutation in histone 3 (H3K27M) redistribute H3K27 acetylation and have increased expression of endogenous retroelements, possibly introducing a vulnerability to XRN1 depletion^33^. Similarly, R882 is a hotspot for mutations in DNMT3A in acute myeloid leukemia and this mutation can lead to expression of immunogenic endogenous retroelements^34^. Rhabdoid tumors could also be a potential target as they are driven by biallelic inactivation of SMARCB1 which also leads to expression of endogenous retroelements^35^.

In addition to inhibiting viral mimicry induction in cells, XRN1 has also been found to be important in the response to viral infections. In combination with viral decapping enzymes, XRN1 can facilitate replication of Vaccinia virus by limiting ISG-inducing dsRNA accumulation^36^. Alternatively, it is possible that degradation of viral RNA can itself pose a threat to viral replication. This is supported by the finding that certain flaviviruses have developed a specific RNA secondary structure that can resist degradation by XRN1^37^. The role of XRN1 in the antiviral response is further highlighted by work by Ran et al. suggesting that XRN1 itself is an ISG whose expression can be induced by IFNα or IFNγ^32^. Together, these results highlight that XRN1 is an important regulator of the antiviral state in the cell, both in exogenous viral infection and endogenous viral mimicry activation.

## Conclusion

Here, we evaluate viral mimicry dependencies across the genome and discover novel targets regulating viral mimicry such as XRN1. XRN1-sensitive cell lines have a higher baseline level of dsRNA and ISGs, while XRN1-resistant cell lines can be made synthetically dependent on XRN1 through viral mimicry-inducing drugs. These results underline the potential for targeting XRN1 in the development of cancer therapeutics against cancers with a high baseline induction in viral mimicry or in combination with viral mimicry-inducing drugs. Interestingly, XRN1 depletion even leads to cell death in cells with ADAR1 overexpression which otherwise can inhibit viral mimicry-dependent cell death, highlighting that XRN1 is a biologically distinct target. Further research is needed to characterize the source of endogenous dsRNA that accumulates upon XRN1 disruption and to establish biomarkers that indicate candidate tumors for XRN1-targeted cancer therapeutics. However, what is clear is that there is a close link between XRN1 and the antiviral state of the cell. Furthermore, these results validate our screening approach for discovery of novel viral mimicry dependencies and suggest that the other hits identified here merit further validation as cancer therapeutic targets.

## Methods

### Primary cell and Cell line Growth conditions

NCI-H747, NCI-H716, COLO678, HCC202 and EFM19 cells were grown in RPMI-1640 medium with 10% (FBS) fetal bovine serum, 2mM glutamine and 1% penicillin/streptomycin. C84 cells were grown in IMDM medium with 10% FBS, 2mM glutamine and 1% Penicillin/Streptomycin. MCF7 cells were grown in DMEM medium with 10% FBS, 1 mM sodium pyruvate, 10 µg/ml human insulin, 0.1mM Non-Essential Amino Acids and 1% penicillin/streptomycin. Patient derived colorectal cells (POP92) were cultured in DMEM/F-12 supplemented with 1% penicillin/streptomycin, 0.1 mM non-essential amino acids, 1 mM sodium pyruvate, N2 supplement, NeuroCult SM1 Neuronal Supplement, 4 μg /ml heparin, 0.2% lipid mixture, 20 ng/ml EGF and 10 ng/ml basic FGF and 1% Penicillin/Streptomycin. All cells were grown according to ATCC recommendations in a humidified tissue culture incubator at 37°C with 5% CO_2_.

### CRISPR Knockout

CRISPR was performed using the Alt-R CRISPR-Cas9 System (Integrated DNA Technologies) as per manufacturer’s protocol. Briefly, the ribonucleoprotein (RNP) complex was assembled by mixing of Alt-R® CRISPR-Cas9 CrRNA with Alt-R® CRISPR-Cas9 tracrRNA and Alt-R® HiFi S.p. Cas9 Nuclease V3. Then cells were transfected with RNP complex by electroporation using pulse code EH115 and Lonza P3 primary cell 4D-Nucleofactor™ X Kit S (Basel, Switzerland).

### In vitro studies and viability assays

The DNA methyltransferase inhibitor 5-AZA-CdR (A3656) was purchased from Sigma Aldrich. Palbociclib isethionate (HY-A0065) was purchased from Med Chem Express. For in vitro experiments, cells were treated with 5-AZA-CdR (300 nM), palbociclib (250 nM) and vehicle (PBS or DMSO) as treatment. Cells were kept in culture for 5 and 7 days for treatment with 5-AZA-CdR and palbociclib respectively. Cell viability was determined using a CellTiter-Glo luminescent cell viability assay (Promega). Data were presented as proliferation present by comparing the treated groups with the vehicle-treated cells.

### Crystal violet staining

Cells were plated in 6-well tissue culture plates for 5 days. Each well was washed twice with ice cold PBS on ice and fixed with 100% ice-cold methanol for about 10 minutes on ice. Then cells were stained with 0.5% crystal violet solution in 25% methanol for 10 min at room temperature, and the crystal violet stain removed by washing in water until the dye comes off.

### Western blot analysis

Protein extracts were obtained from cell pellets after lysis with 100 μL to 200 μL of sodium-dodecyl-sulfate (SDS) lysis buffer (2% SDS, 10% glycerol, 50 mM of Tris HCL) plus protease inhibitor cocktail (Sigma Aldrich, Catalog. No. 11836170001). Cell lysates were centrifuged for 15 min at 4°C, 13000 RPM. 20-40 μg of proteins were mixed with Laemmli (b-mercaptoethanol and bromophenol blue) and denatured for 10 min at 95°C. Cell lysates were loaded onto each lane of SDS-PAGE.

Gel-separated proteins were transferred to a nitrocellulose membrane (Whatman) in a 1X transfer buffer containing 20% methanol, at 100V for 1 hour at 4°C. Membranes were blocked with a solution of in TRIS-buffered saline (TBS: 20mM TRIS/HCl, pH 7.4, 137 mM NaCl, 2.7 mM KCl) plus 0.1% Tween (TBS-T) containing 5% non-fat dried milk. The same milk/TBS-T solution was prepared to dilute primary antibodies, which were incubated for one hour at room temperature or overnight at 4°C. The next day, after 3 washes with 1% TBS-T (each wash 10 minutes), the membrane was incubated with the secondary HRP antibodies, diluted in 5% milk, for 30-60 minutes at room temperature. The membrane was washed again with TBS-T three times and ECL was applied for membrane development.

### Total RNA Extraction and RT-qPCR

For total RNA extraction, 1 mL TRIZOL reagent was added to cell pellets harvested from confluent cells. Cells were gently pipetted up and down for complete cell lysing in TRIZOL. TRIZOL/cell mixture was next transferred into a clean microcentrifuge tube and total RNA was purified using RNeasy Mini Kit (QIAGEN, Valencia, CA) and quantified by spectrophotometer (NanoDrop). Reverse transcription of 1 μg of RNA / sample was performed using SuperScript Vilo IV (Thermo Fisher Scientific) as per manufacturer’s protocol.

5 to 10 ng of cDNA were used to perform quantitative polymerase chain reaction using 1X SsoAdvanced Universal SYBR Green Supermix (Bio-Rad). All RT-qPCR reactions were performed in a CFX Connect Real-Time PCR Detection System (Bio-Rad). Gene expression values were calculated by the ΔCq method, using *RPLP0* as the housekeeping gene, and resulting experimental target values were normalized to the global mean of the control group.

### Dot Blot

Purified RNA was dotted on Hybond N+ membrane (Amerhsam Hybond-N+), dried and autocross linked in a UV stratalinker 2400 (Stratagene) two times. The membrane was then blocked in 5% milk in PBS-T (0.1% Tween-20) for 30 min. Blocking buffer was discarded, and the membrane was incubated with J2 antibody rocking overnight at 4°C. Membrane was washed three times in PBS-T for 10 minutes at room temperature per wash. Membrane was then incubated with secondary goat-anti-mouse HRP antibody (Millipore cat#AP124P) in 5% milk at room temperature for 1 hour. Washed membrane with PBS-T was subjected to chemiluminescent development. Membrane was stained with 0.5% methylene blue in 30% EtOH to visualize the presence of RNA.

### MAVS aggregation assay

Semi-denaturing detergent agarose gel electrophoresis (SDD-AGE) was performed as previously described^38^. Briefly, mitochondria were isolated (Qproteome Mitochondria Isolation Kit, Quiagen) from from patient-derived CRC cells (POP92), resuspended in mitochondria buffer and diluted before loading on a a 1.5% agarose vertical gel at 4°C at a constant voltage of 100 V in running buffer (1x TBE and 0.1% SDS) for 1 hr. Proteins were then transferred into a nitrocellulose membrane, and MAVS protein was detected using anti-MAVS antibody (ab89825, Abcam)

### Immunofluorescence confocal microscopy

Cells were fixed using cold methanol for 15 min at −20°C, washed three times with PBS and incubated with saturation buffer (PBS with 1% BSA) for 1 hour. Cells were stained with primary antibody (anti-dsRNA clone J2, Scicons 10010500) and incubated overnight at 4°C. The next day, cells were washed three times for 10 minutes with PBS and incubated with secondary antibody (anti-mouse IgG (H+L), F(ab’)2 Fragment (Alexa Fluor 647 Conjugate (Cell Signaling Technology, 4410) for 1 hour at room temperature, and washed again three times for 10 minutes with PBS, and incubated with hoechst nuclear stain for 5 minutes. Cells were washed three times with PBS and mounted on a slide with ProLong Gold Antifade Mountant (Thermo Fisher Scientific, P36930). Confocal analysis was performed with a Zeiss LSM70 confocal microscope and images were then analyzed using ImageJ software.

### Linear correlation analysis

CRISPR gene effect and CCLE expression for every gene was downloaded from the DepMap data downloads 22Q2^20^. Only cell lines with both effect and expression were included (n = 1005 cell lines). ISG score was determined by taking the mean of the z-score of the log2(TPM+1) values from the CCLE expression file for the 38 ISGs defined by Liu et al.^22^, see supplementary table 1 for the genelist. For every gene, the linear correlation between the ISG score and gene effect was calculated using the stats.linregress function from the python package scipy. The same approach was used for the other linear correlations in this manuscript.

### STRING gene networks and enrichment analysis

Genelists were submitted to the STRING tool v11.5. Textmining, Experiments, Databases, co-expression, neighborhood, gene fusion and co-occurrence were all used as active interaction sources and medium confidence was used for interaction score. STRING was also used to perform enrichment analysis on the same gene list. The enrichment for the gene ontology “biological process” was downloaded. The 20 terms with the highest strength parameter from STRING were selected and plotted with the ggplot2 package in R.

### Genome-wide search for RNA Editing

The STAR aligner was used to align raw reads to the hg38 human genome^39^, allowing 10 mismatches per aligned read. Duplicate reads were marked and removed, and only unique reads were kept. To avoid bias due to library size, all bam files were down-sampled to 26 million reads. Transcripts were called from each sample using Stringtie^40^, and the called transcripts from all samples were merged using the Stringtie merge mode. To prevent redundant coverage in read positions, the overlap between read mates was clipped using bamUtil clipoverlap^41^, and three base pairs at the start and end of reads were clipped using bamUtil trimbam due to potential errors. Reads aligned to insertions and deletions were discarded.

Variants for each sample in merged transcript regions were called using the bcftools mpileup command^42^, with a minimum read position quality cutoff of Q=20. Indel calling was ignored. Since the RNAseq samples are unstranded, we predicted the strandness of the loci based on several factors such as the strandness of the expressed genes that overlap with the genomic loci.

Filtering process was performed to remove mismatch positions that correspond to common SNPs in the SNP142 common database, which was downloaded from the UCSC genome browser table. A mismatch position is considered present if it has a read depth of at least 5 and if the read depth of the alternative nucleotide is at least 5% of the position read depth and a minimum of 2 reads; otherwise, it is considered absent. To assess editing at Alu elements, edited genomic loci were intersected with the whole genome Alu elements, and only edited loci that overlapped with Alus were kept, while other loci were filtered out.

### Statistical analysis

The number of biological replicates is stated in the figure legends. Unless otherwise stated, statistical analyses were performed using GraphPad prism (v7.0) using Bonferroni Two-way ANOVA and statistical significance was determined at a p value < 0.05.

## Declaration of interests

Daniel De Carvalho reports grants from Princess Margaret Cancer Foundation, the Canadian Institutes of Health Research, and Canada Research Chair during the conduct of the study; grants from Pfizer, and other support from Adela, Inc outside the submitted work. The other authors declare no conflicts of interest.

## Acknowledgments

We thank Catherine O’Brien for providing the POP92 cells.

## References

1. Katze, M.G., He, Y., and Gale, M. (2002). Viruses and interferon: a fight for supremacy. Nat. Rev. Immunol. 2, 675–687. 10.1038/nri888.

2. Chen, R., Ishak, C.A., and De Carvalho, D.D. (2021). Endogenous Retroelements and the Viral Mimicry Response in Cancer Therapy and Cellular Homeostasis. Cancer Discov. 11, 2707–2725. 10.1158/2159-8290.CD-21-0506.

3. Roulois, D., Loo Yau, H., Singhania, R., Wang, Y., Danesh, A., Shen, S.Y., Han, H., Liang, G., Jones, P.A., Pugh, T.J., et al. (2015). DNA-Demethylating Agents Target Colorectal Cancer Cells by Inducing Viral Mimicry by Endogenous Transcripts. Cell 162, 961–973. 10.1016/j.cell.2015.07.056.

4. Chiappinelli, K.B., Strissel, P.L., Desrichard, A., Li, H., Henke, C., Akman, B., Hein, A., Rote, N.S., Cope, L.M., Snyder, A., et al. (2015). Inhibiting DNA Methylation Causes an Interferon Response in Cancer via dsRNA Including Endogenous Retroviruses. Cell 162, 974–986. 10.1016/j.cell.2015.07.011.

5. Mehdipour, P., Marhon, S.A., Ettayebi, I., Chakravarthy, A., Hosseini, A., Wang, Y., de Castro, F.A., Loo Yau, H., Ishak, C., Abelson, S., et al. (2020). Epigenetic therapy induces transcription of inverted SINEs and ADAR1 dependency. Nature 588, 169–173. 10.1038/s41586-020-2844-1.

6. Sheng, W., LaFleur, M.W., Nguyen, T.H., Chen, S., Chakravarthy, A., Conway, J.R., Li, Y., Chen, H., Yang, H., Hsu, P.-H., et al. (2018). LSD1 Ablation Stimulates Anti-tumor Immunity and Enables Checkpoint Blockade. Cell 174, 549‣563.e19. 10.1016/j.cell.2018.05.052.

7. Bowling, E.A., Wang, J.H., Gong, F., Wu, W., Neill, N.J., Kim, I.S., Tyagi, S., Orellana, M., Kurley, S.J., Dominguez-Vidaña, R., et al. (2021). Spliceosome-targeted therapies trigger an antiviral immune response in triple-negative breast cancer. Cell 184, 384‣403.e21. 10.1016/j.cell.2020.12.031.

8. Ishizuka, J.J., Manguso, R.T., Cheruiyot, C.K., Bi, K., Panda, A., Iracheta-Vellve, A., Miller, B.C., Du, P.P., Yates, K.B., Dubrot, J., et al. (2019). Loss of ADAR1 in tumours overcomes resistance to immune checkpoint blockade. Nature 565, 43–48. 10.1038/s41586-018-0768-9.

9. Wells, J.N., and Feschotte, C. (2020). A Field Guide to Eukaryotic Transposable Elements. Annu. Rev. Genet. 54, 539–561. 10.1146/annurev-genet-040620-022145.

10. Griffin, G.K., Wu, J., Iracheta-Vellve, A., Patti, J.C., Hsu, J., Davis, T., Dele-Oni, D., Du, P.P., Halawi, A.G., Ishizuka, J.J., et al. (2021). Epigenetic silencing by SETDB1 suppresses tumour intrinsic immunogenicity. Nature 595, 309–314. 10.1038/s41586-021-03520-4.

11. Wu, Q., Nie, D.Y., Ba-alawi, W., Ji, Y., Zhang, Z., Cruickshank, J., Haight, J., Ciamponi, F.E., Chen, J., Duan, S., et al. (2022). PRMT inhibition induces a viral mimicry response in triple-negative breast cancer. Nat. Chem. Biol., 1–10.

12. Chelmicki, T., Roger, E., Teissandier, A., Dura, M., Bonneville, L., Rucli, S., Dossin, F., Fouassier, C., Lameiras, S., and Bourc‘his, D. (2021). m6A RNA methylation regulates the fate of endogenous retroviruses. Nature 591, 312–316. 10.1038/s41586-020-03135-1.

13. Qiu, W., Zhang, Q., Zhang, R., Lu, Y., Wang, X., Tian, H., Yang, Y., Gu, Z., Gao, Y., Yang, X., et al. (2021). N6-methyladenosine RNA modification suppresses antiviral innate sensing pathways via reshaping double-stranded RNA. Nat. Commun. 12, 1582. 10.1038/s41467-021-21904-y.

14. Gao, Y., Vasic, R., Song, Y., Teng, R., Liu, C., Gbyli, R., Biancon, G., Nelakanti, R., Lobben, K., Kudo, E., et al. (2020). m6A Modification Prevents Formation of Endogenous Double-Stranded RNAs and Deleterious Innate Immune Responses during Hematopoietic Development. Immunity 52, 1007‣1021.e8. 10.1016/j.immuni.2020.05.003.

15. Lindholm, H.T., Chen, R., and De Carvalho, D.D. (2022). Endogenous retroelements as alarms for disruptions to cellular homeostasis. Trends Cancer. 10.1016/j.trecan.2022.09.001.

16. Eisenberg, E., and Levanon, E.Y. (2018). A-to-I RNA editing — immune protector and transcriptome diversifier. Nat. Rev. Genet. 19, 473–490. 10.1038/s41576-018-0006-1.

17. International Human Genome Sequencing Consortium, Whitehead Institute for Biomedical Research, Center for Genome Research:, Lander, E.S., Linton, L.M., Birren, B., Nusbaum, C., Zody, M.C., Baldwin, J., Devon, K., Dewar, K., et al. (2001). Initial sequencing and analysis of the human genome. Nature 409, 860–921. 10.1038/35057062.

18. Ahmad, S., Mu, X., Yang, F., Greenwald, E., Park, J.W., Jacob, E., Zhang, C.-Z., and Hur, S. (2018). Breaching Self-Tolerance to Alu Duplex RNA Underlies MDA5-Mediated Inflammation. Cell 172, 797‣810.e13. 10.1016/j.cell.2017.12.016.

19. Ghandi, M., Huang, F.W., Jané-Valbuena, J., Kryukov, G.V., Lo, C.C., McDonald, E.R., Barretina, J., Gelfand, E.T., Bielski, C.M., Li, H., et al. (2019). Next-generation characterization of the Cancer Cell Line Encyclopedia. Nature 569, 503–508. 10.1038/s41586-019-1186-3.

20. Dempster, J.M., Boyle, I., Vazquez, F., Root, D.E., Boehm, J.S., Hahn, W.C., Tsherniak, A., and McFarland, J.M. (2021). Chronos: a cell population dynamics model of CRISPR experiments that improves inference of gene fitness effects. Genome Biol. 22, 343. 10.1186/s13059-021-02540-7.

21. Jones, C.I., Zabolotskaya, M.V., and Newbury, S.F. (2012). The 5′ → 3′ exoribonuclease XRN1/Pacman and its functions in cellular processes and development. WIREs RNA 3, 455–468. 10.1002/wrna.1109.

22. Liu, H., Golji, J., Brodeur, L.K., Chung, F.S., Chen, J.T., deBeaumont, R.S., Bullock, C.P., Jones, M.D., Kerr, G., Li, L., et al. (2019). Tumor-derived IFN triggers chronic pathway agonism and sensitivity to ADAR loss. Nat. Med. 25, 95–102. 10.1038/s41591-018-0302-5.

23. Gannon, H.S., Zou, T., Kiessling, M.K., Gao, G.F., Cai, D., Choi, P.S., Ivan, A.P., Buchumenski, I., Berger, A.C., Goldstein, J.T., et al. (2018). Identification of ADAR1 adenosine deaminase dependency in a subset of cancer cells. Nat. Commun. 9, 5450. 10.1038/s41467-018-07824-4.

24. Szklarczyk, D., Gable, A.L., Lyon, D., Junge, A., Wyder, S., Huerta-Cepas, J., Simonovic, M., Doncheva, N.T., Morris, J.H., Bork, P., et al. (2019). STRING v11: protein-protein association networks with increased coverage, supporting functional discovery in genome-wide experimental datasets. Nucleic Acids Res. 47, D607–D613. 10.1093/nar/gky1131.

25. O‘Brien, C.A., Kreso, A., Ryan, P., Hermans, K.G., Gibson, L., Wang, Y., Tsatsanis, A., Gallinger, S., and Dick, J.E. (2012). ID1 and ID3 Regulate the Self-Renewal Capacity of Human Colon Cancer-Initiating Cells through p21. Cancer Cell 21, 777–792. 10.1016/j.ccr.2012.04.036.

26. Gao, S., Soares, F., Wang, S., Wong, C.C., Chen, H., Yang, Z., Liu, W., Go, M.Y.Y., Ahmed, M., Zeng, Y., et al. (2021). CRISPR screens identify cholesterol biosynthesis as a therapeutic target on stemness and drug resistance of colon cancer. Oncogene 40, 6601–6613. 10.1038/s41388-021-01882-7.

27. Goel, S., DeCristo, M.J., Watt, A.C., BrinJones, H., Sceneay, J., Li, B.B., Khan, N., Ubellacker, J.M., Xie, S., Metzger-Filho, O., et al. (2017). CDK4/6 inhibition triggers anti-tumor immunity. Nature 548, 471–475. 10.1038/nature23465.

28. Gal-Ben-Ari, S., Barrera, I., Ehrlich, M., and Rosenblum, K. (2019). PKR: A Kinase to Remember. Front. Mol. Neurosci. 11.

29. Li, Y., Banerjee, S., Goldstein, S.A., Dong, B., Gaughan, C., Rath, S., Donovan, J., Korennykh, A., Silverman, R.H., and Weiss, S.R. (2017). Ribonuclease L mediates the cell-lethal phenotype of double-stranded RNA editing enzyme ADAR1 deficiency in a human cell line. eLife 6, e25687. 10.7554/eLife.25687.

30. Chen, Y.G., and Hur, S. (2021). Cellular origins of dsRNA, their recognition and consequences. Nat. Rev. Mol. Cell Biol., 1–16.

31. Pestal, K., Funk, C.C., Snyder, J.M., Price, N.D., Treuting, P.M., and Stetson, D.B. (2015). Isoforms of RNA-Editing Enzyme ADAR1 Independently Control Nucleic Acid Sensor MDA5-Driven Autoimmunity and Multi-organ Development. Immunity 43, 933–944. 10.1016/j.immuni.2015.11.001.

32. Ran, X.-B., Ding, L.-W., Sun, Q.-Y., Yang, H., Said, J.W., Zhentang, L., Madan, V., Dakle, P., Xiao, J.-F., Loh, X., et al. (2023). Targeting RNA exonuclease XRN1 potentiates efficacy of cancer immunotherapy. Cancer Res., CAN-21-3052. 10.1158/0008-5472.CAN-21-3052.

33. Krug, B., De Jay, N., Harutyunyan, A.S., Deshmukh, S., Marchione, D.M., Guilhamon, P., Bertrand, K.C., Mikael, L.G., McConechy, M.K., Chen, C.C.L., et al. (2019). Pervasive H3K27 Acetylation Leads to ERV Expression and a Therapeutic Vulnerability in H3K27M Gliomas. Cancer Cell 35, 782‣797.e8. 10.1016/j.ccell.2019.04.004.

34. Scheller, M., Ludwig, A.K., Göllner, S., Rohde, C., Krämer, S., Stäble, S., Janssen, M., Müller, J.-A., He, L., Bäumer, N., et al. (2021). Hotspot DNMT3A mutations in clonal hematopoiesis and acute myeloid leukemia sensitize cells to azacytidine via viral mimicry response. Nat. Cancer 2, 527–544. 10.1038/s43018-021-00213-9.

35. Leruste, A., Tosello, J., Ramos, R.N., Tauziède-Espariat, A., Brohard, S., Han, Z.-Y., Beccaria, K., Andrianteranagna, M., Caudana, P., Nikolic, J., et al. (2019). Clonally Expanded T Cells Reveal Immunogenicity of Rhabdoid Tumors. Cancer Cell 36, 597‣612.e8. 10.1016/j.ccell.2019.10.008.

36. Burgess, H.M., and Mohr, I. (2015). Cellular 5′-3′ mRNA Exonuclease Xrn1 Controls Double-Stranded RNA Accumulation and Anti-Viral Responses. Cell Host Microbe 17, 332–344. 10.1016/j.chom.2015.02.003.

37. Chapman, E.G., Costantino, D.A., Rabe, J.L., Moon, S.L., Wilusz, J., Nix, J.C., and Kieft, J.S. (2014). The Structural Basis of Pathogenic Subgenomic Flavivirus RNA (sfRNA) Production. Science 344, 307–310. 10.1126/science.1250897.

38. Ettayebi, I., Yau, H.L., and De Carvalho, D.D. (2019). Methods to detect endogenous dsRNA induction and recognition. In Methods in Enzymology (Elsevier), pp. 35–51. 10.1016/bs.mie.2019.07.002.

39. Dobin, A., Davis, C.A., Schlesinger, F., Drenkow, J., Zaleski, C., Jha, S., Batut, P., Chaisson, M., and Gingeras, T.R. (2013). STAR: ultrafast universal RNA-seq aligner. Bioinformatics 29, 15–21. 10.1093/bioinformatics/bts635.

40. Pertea, M., Pertea, G.M., Antonescu, C.M., Chang, T.-C., Mendell, J.T., and Salzberg, S.L. (2015). StringTie enables improved reconstruction of a transcriptome from RNA-seq reads. Nat. Biotechnol. 33, 290–295. 10.1038/nbt.3122.

41. Jun, G., Wing, M.K., Abecasis, G.R., and Kang, H.M. (2015). An efficient and scalable analysis framework for variant extraction and refinement from population-scale DNA sequence data. Genome Res. 25, 918–925. 10.1101/gr.176552.114.

42. Narasimhan, V., Danecek, P., Scally, A., Xue, Y., Tyler-Smith, C., and Durbin, R. (2016). BCFtools/RoH: a hidden Markov model approach for detecting autozygosity from next-generation sequencing data. Bioinformatics 32, 1749–1751. 10.1093/bioinformatics/btw044.

