## Supplementary figures for "Retroelement decay by the exonuclease XRN1 is a viral mimicry dependency in cancer"

**Supplementary file**

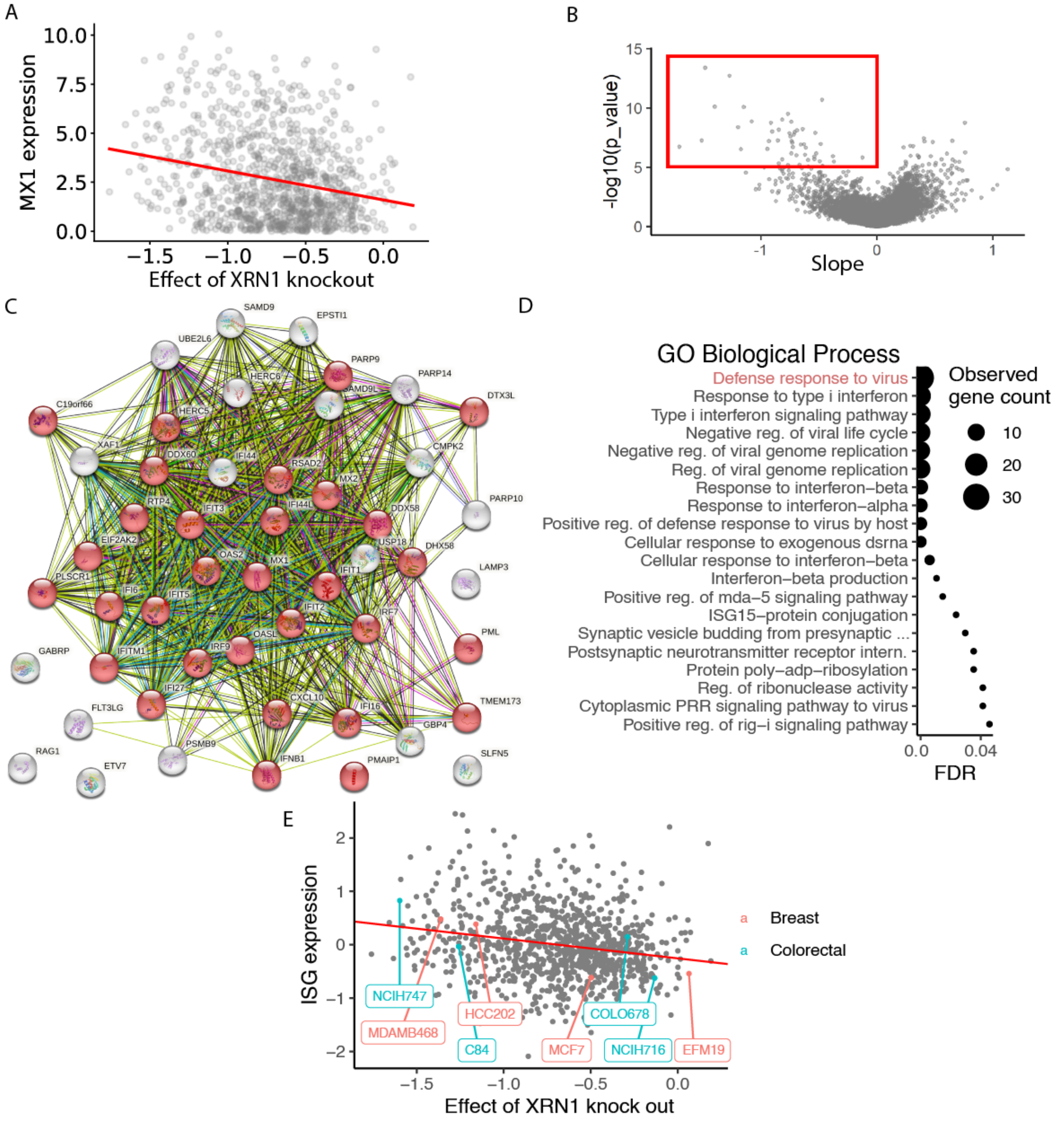


**Figure S1.** Correlation between effect of XRN1 knock out on viability and ISG expression. (A) Example of linear correlation analysis between gene expression from CCLE and effect on viability by knockout of XRN1 from the CRISPR DepMap dataset from 1005 cell lines. Here shown for the gene MX1. (B) Slope and p-value from linear correlation analysis as done in A. Red box shows the top 50 most significant genes with negative slope. (C) Network of the top 50 genes most significantly correlating with effect on viability from XRN1 knockout. Made with the STRING tool. (D) The 20 terms with the highest strength of the biological process GO terms according to the STRING tool. (E) Linear correlation between mean scaled ISG expression in the CCLE dataset of 38 ISGs defined by Liu et al and effect on viability after knocking out XRN1 from the DepMap dataset. Each dot is a cell line and here we have highlighted cell lines studied further in this project.

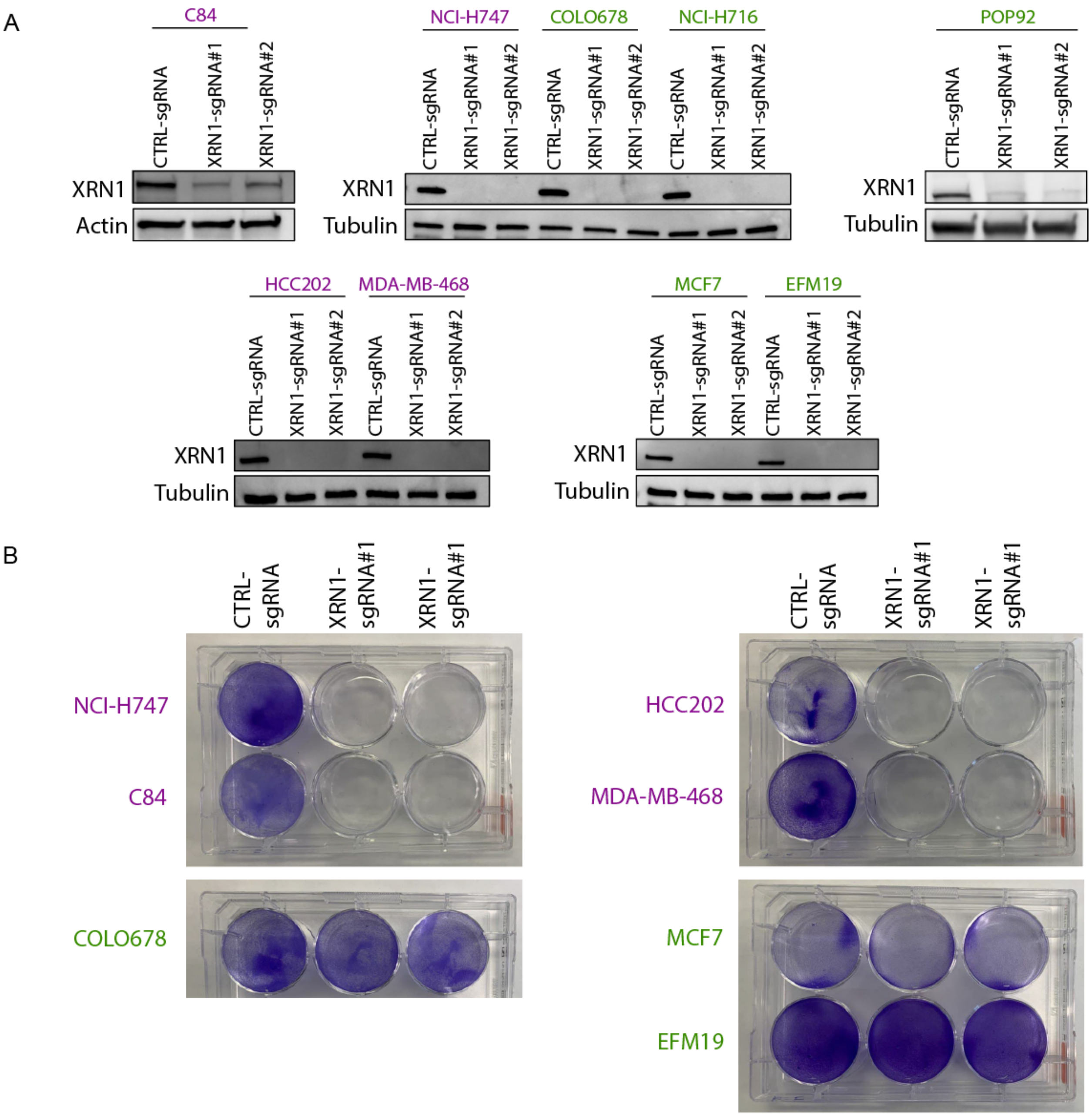


**Figure S2.** Viability of cell lines after XRN1 knockout. (A) Western blot of XRN1 protein level in indicated cell lines. XRN1-sensitive cell lines are colored purple and XRN1-resistant cell lines are colored green. Two independent clones were used for XRN1-KO. Tubulin or actin was used as a loading control. (B) Cell viability of control or XRN1-KO cells assessed by crystal violet staining.

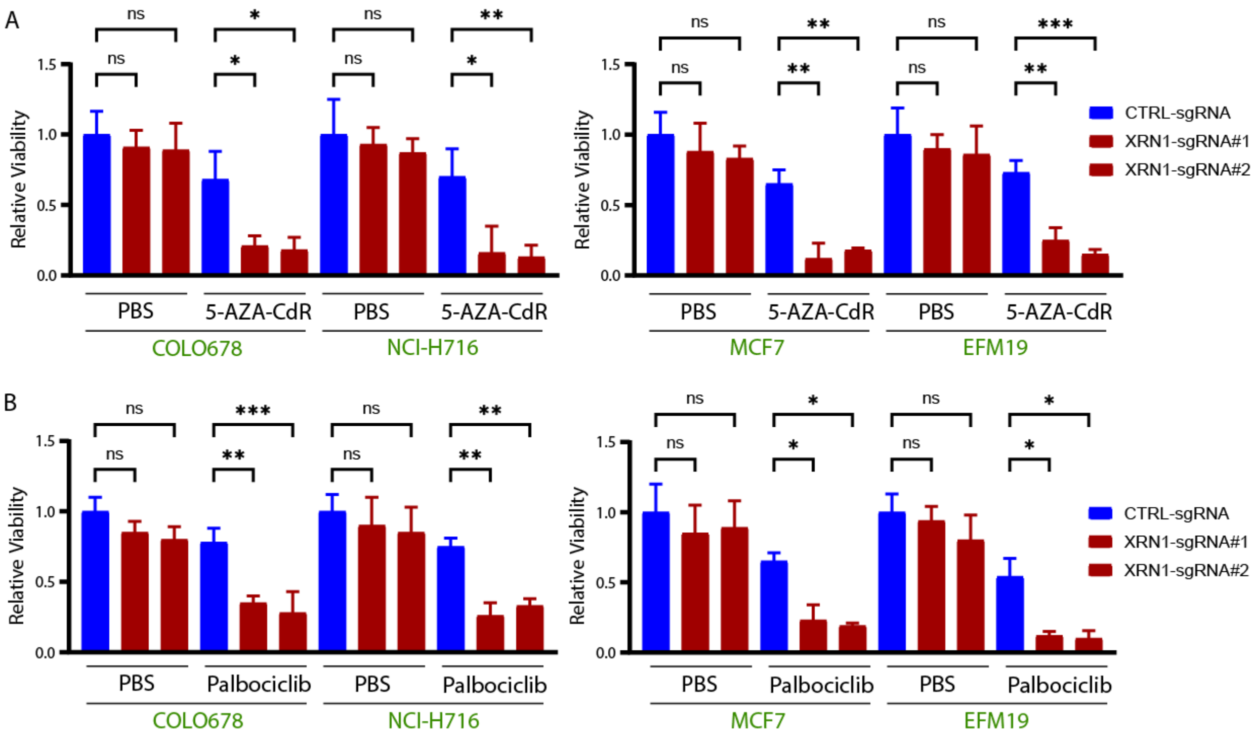


**Figure S3.** AZA and Palbociclib sensitizes XRN1-resistant cell lines to XRN1 knockout. (A) Cell viability was assessed by celltiter glo in CTRL and XRN1 KO Colorectal and breast cancer cells treated with 5-AZA-CdR. (B) Cell viability was assessed by celltiter glo in CTRL and XRN1 KO Colorectal and breast cancer cells treated with Palbociclib. Data presented as the mean of triplicates ± SD.

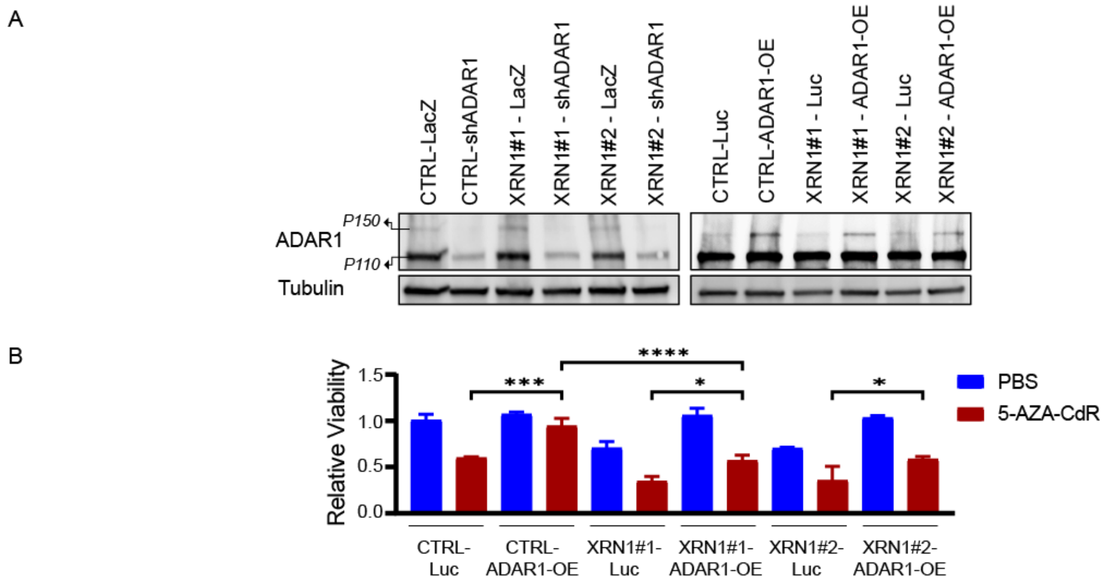


**Figure S4**. Depletion of XRN1 and over expression of ADAR1 in POP92 cells. A) Western blot of ADAR1 in POP92 cells with ADAR1 and XRN1 knockout and ADAR1 overexpression. Tubulin is loading control. sgRNA is used to knockout XRN1, #1 and #2 represents different sgRNA targeting XRN1. B) Extension of figure 5D including additional comparisons. Cell viability in POP92 cells with indicated knockout or overexpression determined with Cell Titer Glo assay. Data presented as the mean of triplicates ± SD.

### References


1. Liu, H., Golji, J., Brodeur, L.K., Chung, F.S., Chen, J.T., deBeaumont, R.S., Bullock, C.P., Jones, M.D., Kerr, G., Li, L., et al. (2019). Tumor-derived IFN triggers chronic pathway agonism and sensitivity to ADAR loss. Nat. Med. *25*, 95–102. 10.1038/s41591-018-0302-5.
